# Genetic modulation of mitochondrial NAD+ regeneration does not prevent dopaminergic neuron dysfunction caused by mitochondrial complex I impairment

**DOI:** 10.1101/2025.06.07.658365

**Authors:** Karis B. D’Alessandro, Enrico Zampese, Jenna L. E. Blum, Britta Kuusik, Alec Palmiotti, Shawn M. Davidson, Colleen R. Reczek, D. James Surmeier, Navdeep S. Chandel

## Abstract

Dysfunction of mitochondrial complex I (MCI) has been implicated in the degeneration of dopaminergic neurons in Parkinson’s disease. Here, we report the effect of expressing MitoLbNOX, a mitochondrial-targeted version of the bacterial enzyme LbNOX, which increases regeneration of NAD+ in the mitochondria to maintain the NAD+/NADH ratio, in dopaminergic neurons with impaired MCI (MCI-Park mice). MitoLbNOX expression did not ameliorate the cellular or behavioral deficits observed in MCI-Park mice, suggesting that alteration of the mitochondrial NAD+/NADH ratio alone is not sufficient to compensate for loss of MCI function in dopaminergic neurons.

## Introduction

Dopaminergic neurons are key modulators of brain function. They are implicated in the regulation of mood, reward circuits, motivation, and voluntary movements.^1^ Previous research has suggested that dopaminergic neurons are more metabolically demanding than other neuronal subtypes, and thus more vulnerable to disruptions in mitochondrial function.^2^

Mitochondrial complex I (MCI) of the electron transport chain has been implicated in the pathogenesis of Parkinson’s disease (PD) through various known genetic and environmental contributors to the disease.^3,4^ MCI serves three primary functions: proton pumping, production of reactive oxygen species (ROS), and regeneration of NAD+ from NADH.^5^ Recently, a mouse model with conditional loss of MCI in dopaminergic neurons was developed (i.e., MCI-Park mice).^6^ This model crosses *Ndufs2*-floxed mice with a dopaminergic neuron-specific Cre (DAT-Cre). NDUFS2 is a catalytic subunit necessary for the function of the forty-five subunit mammalian MCI.^7^ MCI-Park mice develop a progressive, levodopa-responsive Parkinsonism, characterized by early loss of tyrosine hydroxylase expression in the substantia nigra pars compacta (SNc) and progressive motor disability. In these MCI-Park mice, fine motor function is disrupted at postnatal day 30 (P30), while gross motor dysfunction begins around P60. These motor deficits are paralleled by a progressive loss of tyrosine hydroxylase expression, which starts in axons and later becomes evident in cell bodies. This phenotypic down-regulation is followed by frank neurodegeneration, weight loss, and mortality.

Previous work has attempted to mitigate the loss of MCI function, notably through expression of the *Saccharomyces cerevisiae* protein NDI1. NDI1 expression regenerates NAD+ from NADH and donates two electrons to the CoQ pool of the electron transport chain. NDI1 expression effectively restores both NAD+ regeneration and electron transport chain function.^8–10^ Importantly, NDI1 cannot pump protons or generate superoxide, and thus, NDI1 does not directly contribute to ATP generation or ROS production, respectively. Nevertheless, previous studies have shown that NDI1 expression protects dopaminergic neurons in toxin-based models of PD,^11–14^ and our ongoing work indicates that NDI1 expression protects dopaminergic neurons in MCI-Park mice.^15^ The impact of NDI1 expression on dopaminergic neurons is attributable either to the regeneration of NAD+ from NADH or to its ability to support electron transport. In support of the former possibility, NAD+ supplementation via administration of precursor nicotinamide riboside protects dopaminergic neurons, suggesting that alteration of the NAD+/NADH ratio may be sufficient to ameliorate neurodegeneration.^16^ Phase I clinical trials of NAD+ precursor supplementation were conducted in patients with PD.^17^

To test the hypothesis that NAD+ regeneration is sufficient to prevent neurodegeneration of dopaminergic neurons due to MCI loss, an NADH oxidase from *Lactobacillus brevis* that regenerates NAD+ from NADH and is localized to the mitochondrial matrix (MitoLbNOX) was expressed in the MCI-Park mouse.^18^ Unlike NDI1, MitoLbNOX does not restore electron transport chain function, but does regenerate mitochondrial NAD+ from NADH.^19^ Contrary to our hypothesis, the expression of MitoLbNOX did not prevent the loss of dopaminergic neurons and Parkinsonism in MCI-Park mice.

## Results

To test whether increasing mitochondrial NAD+ regeneration alone in dopaminergic neurons in MCI-Park mice would ameliorate their Parkinsonian phenotype, we designed a targeting construct containing *Rosa26* homology arms and a Lox-STOP-Lox (LSL) cassette upstream of the MitoLbNOX gene. Thus, MitoLbNOX is only expressed following Cre-mediated removal of the LSL cassette. We crossed MitoLbNOX-LSL mice with MCI-Park mice to induce MitoLbNOX expression and NDUFS2 loss specifically in dopaminergic neurons (Fig. 1a). MCI-Park mice have been reported to have a median survival of ∼21.5 weeks.^20^ We found MCI-Park mice to have a median survival of 28 weeks. Expression of one allele of MitoLbNOX in MCI-Park mice increased their median survival to 44 weeks, but this effect was not statistically significant (Fig. 1b). Expression of two alleles of MitoLbNOX in MCI-Park mice did not significantly alter the survival of mice compared to expression of one allele. Mice expressing MitoLbNOX without the concomitant loss of NDUFS2 (DAT-Cre + MitoLbNOX LSL mice) had a similar median survival as control DAT-Cre mice (Fig. 1c).

**Figure 1.**
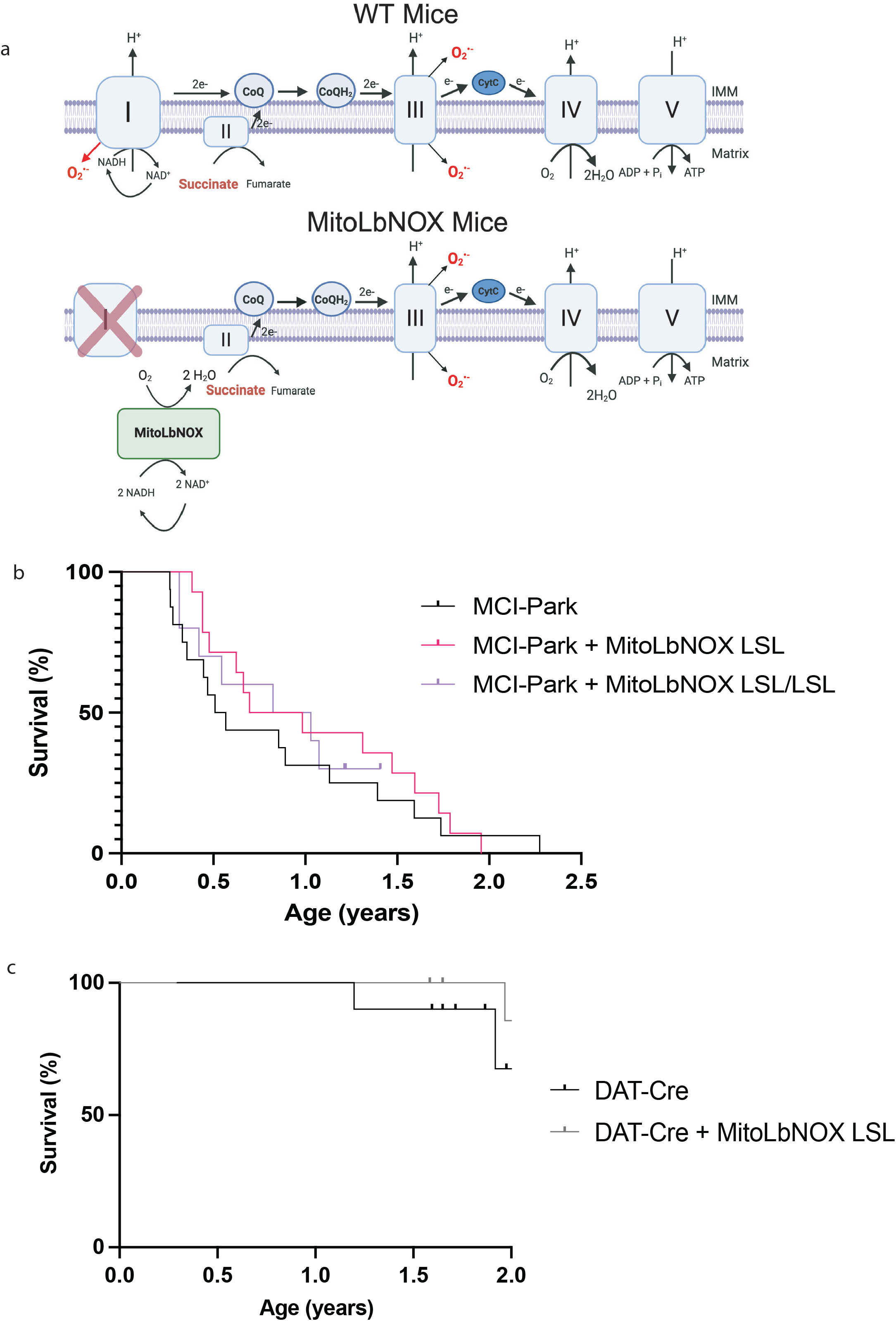
MitoLbNOX expression does not restore lifespan of MCI-Park mice. a) MitoLbNOX regenerates NAD+ from NADH in the mitochondria. In a setting with mitochondrial complex I deficiency, MitoLbNOX is unable to restore proton pumping, superoxide production, or the transfer electrons to the downstream electron transport chain complexes. In MCI-Park + MitoLbNOX LSL mice, Exon 2 of the Ndufs2 gene is flanked by LoxP sites. The MitoLbNOX gene is downstream of a Lox-STOP-Lox cassette. Cre recombinase expression under the control of the dopamine transporter (DAT) reporter allows for expression of MitoLbNOX and loss of the catalytic subunit of mitochondrial complex I NDUFS2 specifically in the dopaminergic neurons of MCI-Park + MitoLbNOX LSL mice. Created in BioRender. Chandel, N. (2025) https://BioRender.com/7wy3r6d b) MitoLbNOX expression moderately extends the lifespan of MCI-Park mice (n=10-16 mice, Log-Rank (Mantel-Cox) test not significant). Marked points represent mice that had not yet met endpoint criteria at the end of the study. c) MitoLbNOX expression in dopamine neurons in DAT-Cre positive mice with a functional endogenous mitochondrial complex I does not alter lifespan (n=9-10 mice, Log-Rank (Mantel-Cox) test not significant). Marked points represent mice that had not yet met endpoint criteria at the end of the study.

Next, we performed an open field assay to measure the motor phenotype at 30, 60, and 100 days of age. In agreement with previous work,^6^ both distance and velocity traveled were significantly lower in MCI-Park mice compared to DAT-Cre controls at P30, P60, and P100 (Fig. 2a-c, Supp. Fig. 1a-d). MitoLbNOX expression, one or two alleles, in MCI-Park mice did not improve open field behavior at any timepoint (Fig. 2a-c, Supp. Fig. 1a-d). Rotarod testing was also performed to further quantify motor phenotype. MCI-Park mice exhibit a significantly shorter fall latency compared to DAT-Cre mice at all timepoints (Fig. 2d, Supp. Fig. 1e-f), consistent with previous findings that show fine motor function is disrupted in MCI-Park mice as early as P30.^6^ Expression of MitoLbNOX, one or two alleles, in MCI-Park mice did not improve performance on the rotarod task at any time point (Fig. 2d, Supp. Fig. 1e-f). Importantly, expression of MitoLbNOX in DAT-Cre mice did not negatively affect motor phenotype (Supp. Fig. 2a-i).

**Figure 2.**
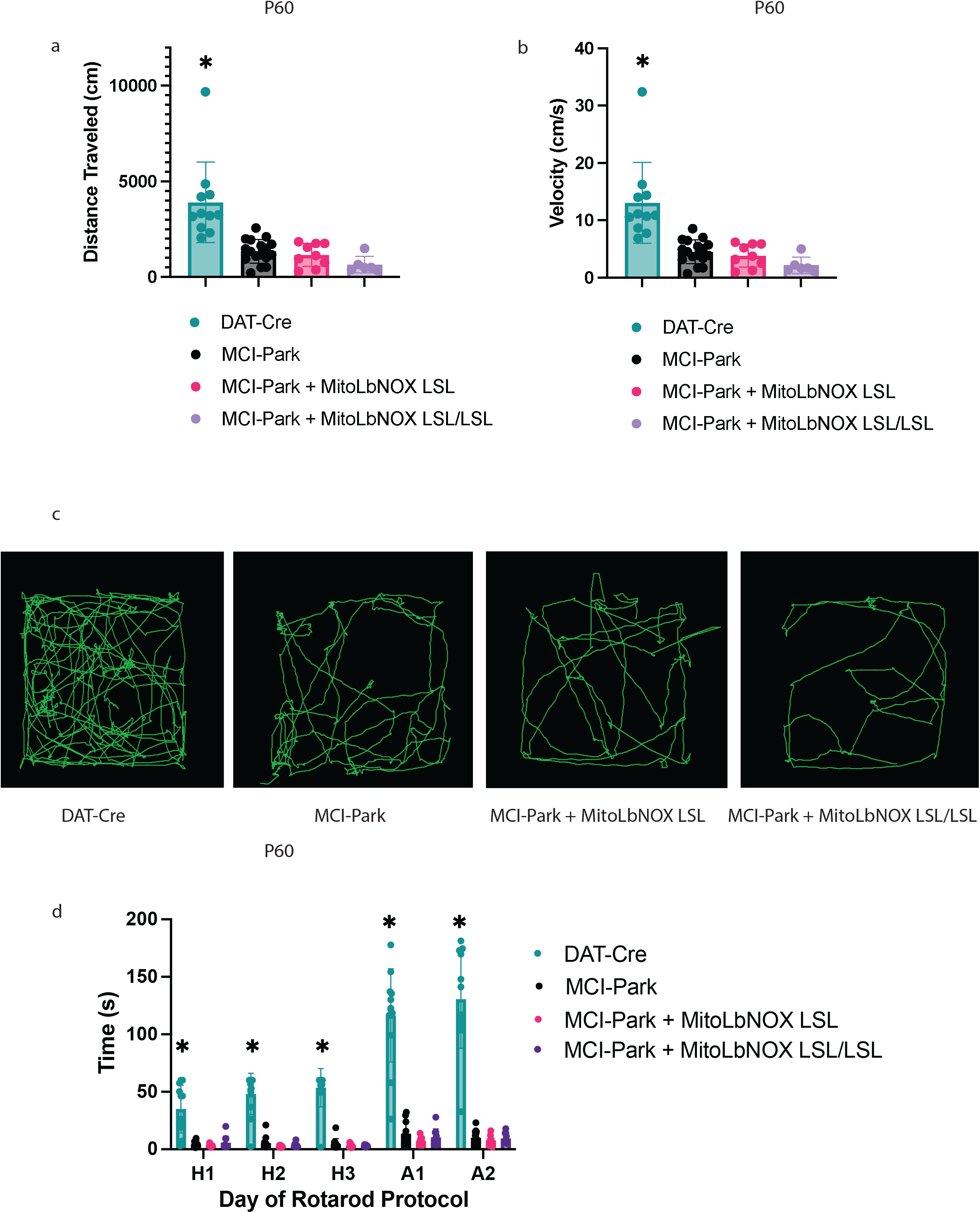
MitoLbNOX expression does not restore motor function in MCI-Park mice. a) MitoLbNOX did not improve distance traveled in an open field test at P60 (n=6-17 mice, Welch’s t-test p<0.01 for DAT-Cre mice compared to each other group, not significant for comparisons between MCI-Park, MCI-Park + MitoLbNOX, and MCI-Park + MitoLbNOX LSL/LSL). b) MitoLbNOX did not improve velocity traveled in an open field test at P60 (n=6-17 mice, Welch’s t-test p<0.01 for DAT-Cre mice compared to each other group, not significant for comparisons between MCI-Park, MCI-Park + MitoLbNOX, and MCI-Park + MitoLbNOX LSL/LSL). c) Representative tracings of open field data. d) MitoLbNOX expression does improve rotarod performance in MCI-Park mice (n=8-13, multiple unpaired t-test p<0.001 for DAT-Cre mice compared to each other group on all days, not significant for comparisons between MCI-Park, MCI-Park + MitoLbNOX, and MCI-Park + MitoLbNOX LSL/LSL on any days). Days H1-H3 represent a three-day habituation period with a maximum time of 60 seconds at a constant speed. Days A1-A2 represent a two-day acceleration period with a maximum time of 300 seconds at an accelerating speed.

To determine whether MitoLbNOX expression in MCI-Park mice induces metabolic changes, we performed bulk metabolomics via liquid chromatography-mass spectrometry (LC-MS) on microdissected SNc samples at P45-60. Because the number of copies of the MitoLbNOX-LSL allele did not have a significant impact on phenotypic output, we performed our bulk metabolomics study on mice containing one MitoLbNOX-LSL allele. Partial Least Squares Discriminant Analysis (PLS-DA) revealed that DAT-Cre, MCI-Park, and MCI-Park + MitoLbNOX LSL mice were distinct; however, MCI-Park and MCI-Park + MitoLbNOX LSL mice were closer to each other than to DAT-Cre mice (Fig. 3a).

**Figure 3.**
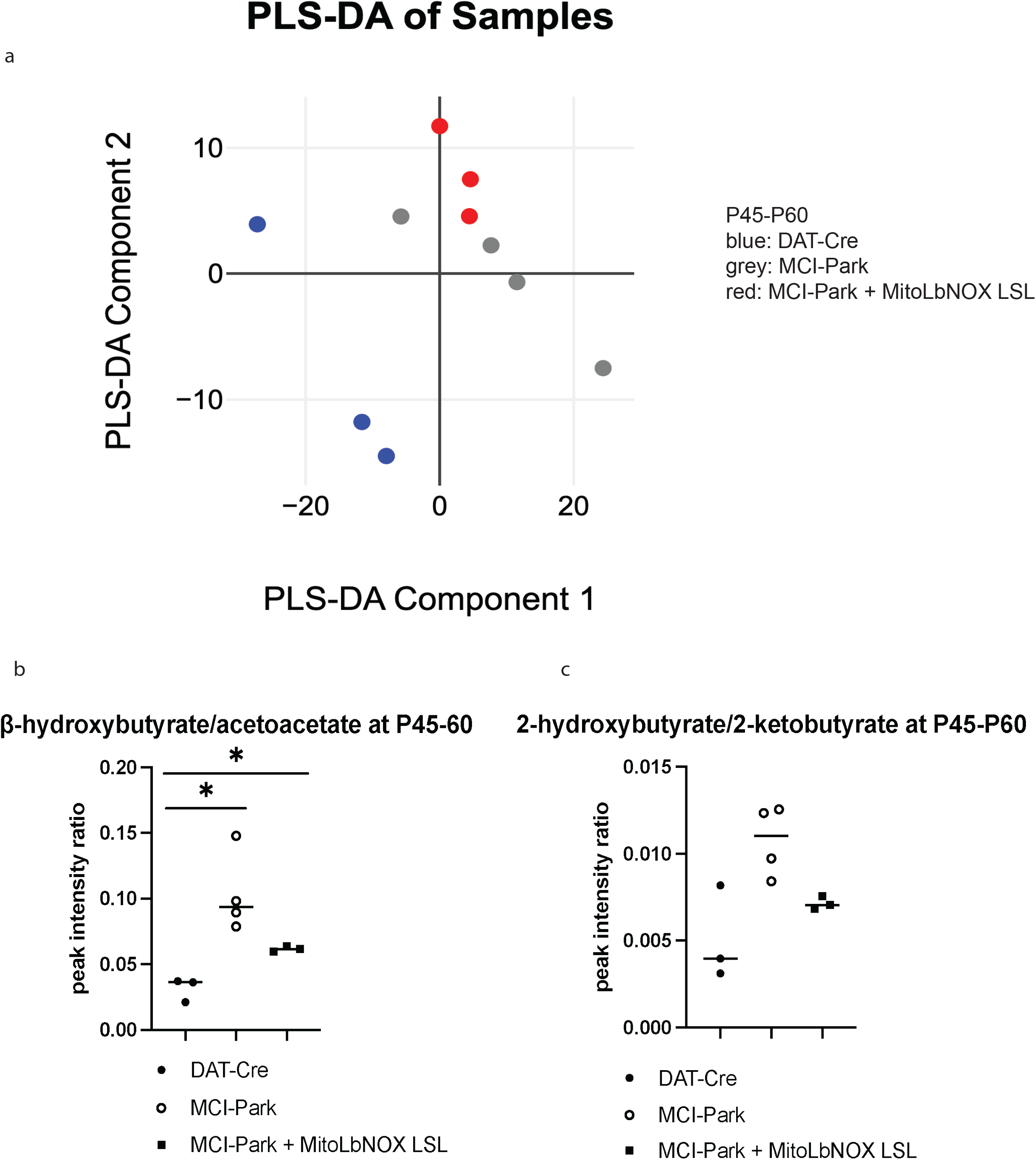
MitoLbNOX expression lowers the mitochondrial NADH/NAD+ ratio in MCI-Park mice. a) Partial least squares discriminant analysis (PLS-DA) shows distinctive clustering between the genotypes (n=3-4 mice). b) β-hydroxybutyrate/acetoacetate ratio is elevated in MCI-Park mice compared to DAT-Cre mice. MitoLbNOX expression in MCI-Park mice diminishes this ratio, though not significantly (n=3-4 mice, Welch’s t-test p=0.0135 for DAT-Cre compared to MCI-Park, p=0.0237 for DAT-Cre compared to MCI-Park + MitoLbNOX LSL, not significant (p=0.0709) for MCI-Park compared to MCI-Park + MitoLbNOX LSL). c) 2-hydroxybutyrate/2-ketobutyrate ratio is elevated in MCI-Park mice compared to DAT-Cre mice. MitoLbNOX expression in MCI-Park mice diminishes this ratio, though not significantly (n=3-4 mice, Welch’s t-test not significant).

Next, we examined NAD+-associated metabolites to determine if expression of MitoLbNOX in our MCI-Park mice alters the NADH/NAD+ ratio. β-hydroxybutyrate and acetoacetate are ketone bodies that serve as fuel sources in brain mitochondria.^21^ Acetoacetate conversion to β-hydroxybutyrate is coupled to a reduction of NADH to produce NAD+. Thus, an increase in the β-hydroxybutyrate/acetoacetate ratio reflects a high mitochondrial NADH/NAD+ ratio. Similarly, the conversion of 2-ketobutyrate to 2-hydroxybutyrate is coupled to the NADH/NAD+ ratio. An increase in the 2-hydroxybutyrate/2-ketobutyrate ratio also suggests an elevated mitochondrial NADH/NAD+ ratio. Examination of these metabolite ratios have previously been used as a proxy for mitochondrial NADH/NAD+ ratio in other models, including those caused by loss-of-function mutations in MCI subunits.^22–24^ Both the β-hydroxybutyrate/acetoacetate and the 2-hydroxybutyrate/2-ketobutyrate ratios were elevated in the MCI-Park mice compared to the DAT-Cre mice, which was reduced by expression of MitoLbNOX (Fig. 3b and c). These data indicate that MitoLbNOX was functional and able to lower the mitochondrial NADH/NAD+ ratio observed in MCI-Park mice.

To functionally determine if these motor impairments were due to the loss of tyrosine hydroxylase (TH), we performed immunocytochemistry. At P45-60, the MCI-Park mice had significantly fewer TH+ SNc neurons than DAT-Cre mice (Figure 4a-b). Expression of one allele of MitoLbNOX in MCI-Park mice was not sufficient to restore the number of TH+ neurons (Figure 4a-b). TH+ levels in the ventral tegmental area (VTA) of MCI-Park and MCI-Park + MitoLbNOX LSL mice were not significantly lower than those of DAT-Cre mice at this time point (Fig. 4a, c).

**Figure 4.**
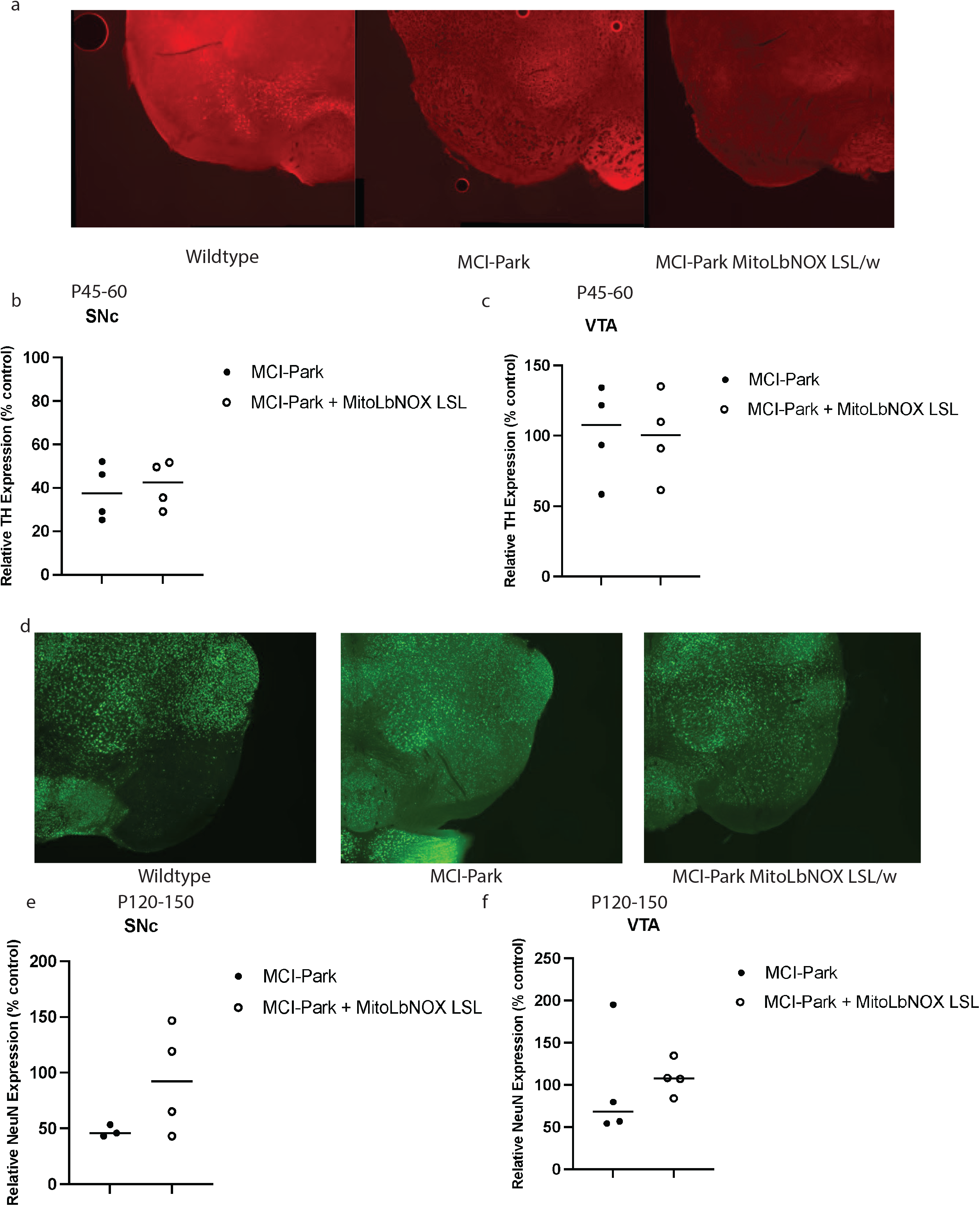
MitoLbNOX expression does not restore tyrosine hydroxylase (TH) expression or prevent cell death. a) Representative TH+ staining shows significant loss at P45-60 in MCI-Park mice that is not restored by MitoLbNOX expression. b) MCI-Park mice show a significant reduction in TH+ expression in the substantia nigra pars compacta (SNc) at P45-60 as compared to DAT-Cre control mice that is not restored with MitoLbNOX expression (n=4, Welch’s t-test p=0.0470 for DAT-Cre compared to MCI-Park, not significant for any other comparisons). c) MCI-Park and MCI-Park + MitoLbNOX LSL mice do not show a significant reduction in TH+ expression in the ventral tegmental area (VTA) at P45-60 as compared to DAT-Cre control mice (n=4, Welch’s t-test not significant). d) Representative NeuN staining, a measure of cell death, shows a loss at P120-150 in MCI-Park mice that is not restored by MitoLbNOX expression. e) MCI-Park mice show a significant reduction in NeuN expression in the SNc at P120-150 as compared to DAT-Cre control mice that is not restored with MitoLbNOX expression (n=4, Welch’s t-test p=0.0089 for DAT-Cre compared to MCI-Park, not significant for any other comparisons). f) MCI-Park and MCI-Park + MitoLbNOX LSL mice do not show a significant reduction in NeuN expression in the VTA at P120-150 as compared to DAT-Cre control mice (n=4, Welch’s t-test not significant between any groups).

As MCI-Park mice age, the number of SNc dopaminergic neurons decreases.^6^ We performed neuronal nuclei (NeuN) immunostaining in the SNc of P120-150 aged mice to quantify neuronal death. MCI-Park mice expressing MitoLbNOX had an increased number of NeuN-positive neurons in the SNc (i.e., less neuronal cell death) compared to MCI-Park mice; however, this difference was not statistically significant (Fig. 4d-e). The number of NeuN-positive neurons in the VTA was also increased, albeit not significantly, at this age in MCI-Park mice expressing MitoLbNOX (Fig. 4d, f).

## Discussion

Our findings suggest that enhancing mitochondrial NAD^+^ regeneration to maintain the NAD^+^/NADH balance is insufficient to prevent the loss of dopaminergic neurons and onset of Parkinsonian symptoms in MCI-Park mice. MitoLbNOX expression did not prevent the onset of Parkinsonism, downregulation in the dopaminergic neuron phenotype, or eventual neurodegeneration in MCI-Park mice. The modest extension in lifespan and increased NeuN staining observed in P120-150 mice suggest that MitoLbNOX might slightly delay cell loss, although these changes did not reach statistical significance. Importantly, while behavioral and lifespan assays were adequately powered, the samples sizes for immunocytochemistry were limited, which may have impacted statistical significance.

One important limitation of this study was our inability to directly measure the NADH/NAD+ ratio in SNc dopaminergic neurons. This prevented an accurate assessment of the extent to which the NADH/NAD+ ratio was restored by MitoLbNOX in MCI-Park mice. Additionally, MitoLbNOX may be unable to sustain mitochondrial NAD+ regeneration at levels comparable to those of endogenous MCI in dopaminergic neurons *in vivo*.

## Methods

### Animals

Ndufs2 floxed mice were provided by J. López-Barneo (Universidad de Sevilla, Spain). DAT-Cre mice (B6.SJL-*Slc6a3*^*tm1*.*1(cre)Bkmn*^/J) were provided by D. James Surmeier (Northwestern University). MitoLbNOX mice were generated at Northwestern University’s Transgenic and Targeted Mutagenesis Laboratory. The MitoLbNOX gene was cloned into our previously published targeting construct.^10,25^

For Kaplan-Meier survival curve analysis, mice were considered events (deaths) if they lost >10% of their maximum body weight, exhibited decreased responsiveness, and appeared lethargic. Data was censored for animals euthanized due to unrelated health issues and for experimental analysis prior to reaching a terminal state.

Mice were maintained in Northwestern University’s Center for Comparative Medicine (CCM) in individually ventilated microisolator cages and were provided ad libitum access to standard rodent chow (Envigo/Teklad LM-485), automatic water dispenser, and long sipper tube water bottles. Mice were additionally provided with moist chow on the cage bed beginning at P30. Housing conditions included a 12-hour light/dark cycle, humidity of 30-70%, ambient temperature of 72 +/-2 degrees Fahrenheit, and biweekly cage changes, in accordance with CCM and Northwestern University’s Institutional Animal Care and Use Committee’s (IACUC’s) guidelines. Mice were monitored three times weekly and weighed once weekly to ensure animal welfare. Males and females were used for all studies. Subgroup analysis did not reveal sex-specific differences on any of the quantified metrics.

All animal procedures were reviewed and approved by the IACUC at Northwestern University.

### Open Field Test

Mice were placed in an open field chamber (56 cm x 56 cm) within a soundproof box and spontaneous activity was recorded for 300 seconds. Distance traveled from time 0-299 seconds and average velocity traveled from 0-299 seconds were recorded and analyzed using the Limelight (v5) software. Open field testing was performed at +/-5 days from each timepoint depending on instrument availability. Open field testing was performed using equipment at Northwestern University’s Behavioral Phenotyping Core.

### Rotarod Test

For rotarod testing, mice were placed onto a spinning cylindrical rod device (RotaRod) upon which constant forward movement was required to prevent falling. The maximum length of time spent on the RotaRod based on the day of protocol and latency to fall was recorded using Rod software. The rotarod protocol involved 5 consecutive days of testing. For the first three days (habituation), mice were placed on the rod that was rotating a constant speed of 12 rotations/min for a maximum of 60 seconds. For the last two days of the protocol (acceleration), mice were placed on the rod which was accelerating at a constant rate from 4 to 40 rotations/min for a maximum of 300 seconds. Testing was repeated four times each day per mouse with a 5–10-minute rest period between trials. A trial was repeated if a mouse had a latency to fall of less than one second. Rotarod testing was performed at +/-5 days from each timepoint depending on instrument availability. Rotarod testing was performed using equipment at Northwestern University’s Behavioral Phenotyping Core.

### LC-MS

SNc tissue was micro-dissected, snap-frozen on dry ice, and stored at −80 °C until extraction. Polar metabolites were extracted on ice with 40:40:20 acetonitrile:methanol:HPLC-grade water with 0.1M formic acid (-20 °C) containing 200 ng/mL isotope-labeled internal standards (thymine-d_4_ and inosine-^15^N_4_ for negative ion mode; valine-d_8_ and phenylalanine-d_8_ for positive ion mode).Tissue was mechanically dissociated via pipetting up and down, vortexed briefly, and incubated on ice for 10 mins. Another extraction buffer was then added of HPLC-grade water with 15% w/v ammonium bicarbonate. Lysates were cleared by centrifugation (14,000 x g, 25 min, 4°C). 100 μL of the clarified extracts were transferred to autosampler vials for LC-MS.

Chromatographic separation was achieved on an XBridge BEH Amide HILIC column (2.1 × 150 mm, 2.5 μm; Waters) using an UltiMate 3000 UHPLC (Thermo Fisher Scientific). Mobile phase A was 95 % H_2_O/5 % acetonitrile with 20 mM ammonium acetate and 20 mM ammonium hydroxide, adjusted to pH 9.4; mobile phase B was acetonitrile. The gradient (0.3 mL/min, 25°C column temperature, 10 μL injection) was: 0-2 min 90% B; 2-3 min 75% B; 3-7 min 75% B; 7-8 min 70% B; 8-10 min 70→50% B; 10-12 min 50% B; 12-13 min 25% B; 13-16 min 25→0% B; 16-21 min 0% B; 21-25 min re-equilibration at 90% B.

Mass spectra were acquired on an Orbitrap Exploris 240 (Thermo Fisher Scientific) operated in polarity-switching full-scan mode (m/z 70-1,000 Da, resolution 60,000 FWHM at m/z 200; spray voltage 2.8 kV for negative ionization, 3.2 kV for positive ionization; capillary temperature 320°C; source temperature 30°C; sheath/aux/sweep gas 35/10/0 arb).

Raw files were processed in El-MAVEN (Elucidata.io). Peak areas were normalized to the corresponding isotope-labeled internal standard whenever available; metabolites lacking a dedicated standard were normalized to total ion current (TIC). Normalized data were exported for downstream statistical analysis.

### Immunocytochemistry

Mice were perfused with ice cold PBS and 4% paraformaldehyde (PFA), and brains were dissected. Following dissection, brains were stored in 4% PFA solution for 12-16 hours before being placed in 15% sucrose diluted in Dulbecco’s Phosphate-Buffered Saline (DPBS) for 12-16 hours. Brains were then stored in 30% sucrose until sectioning. Brains were sectioned coronally into 30 μm thick sections by vibratome (Leica VT1200S) and stored in 0.1% sodium azide diluted in DPBS before staining.

For tyrosine hydroxylase (TH) staining, sections containing the region of interest were selected and rinsed in DPBS followed by washes in PBS with Triton. Sections were then incubated in a PBS with Triton solution containing donkey serum for 30 minutes at room temperature. Sections were incubated overnight at 4°C in Tyrosine Hydroxylase Antibody (Immunostar, 22941) at a 1:1000 ratio. Following this incubation in primary antibody, sections were again washed in PBS with Triton and then incubated at room temperature in Goat anti-Mouse IgG (H+L) Cross-Adsorbed Secondary Antibody, Alexa Fluor™ 594 (ThermoFisher Scientific, A-11005) for 60 minutes at a 1:400 ratio. Sections were then rinsed with PBS with Triton, then DPBS before mounting on slides (Fisher Scientific, 12-550-15). Once dry, sections were coverslipped using ProLong™ Gold Antifade Mountant with DNA Stain DAPI (ThermoFisher Scientific, P36935). Slides were stored at room temperature in dark conditions overnight to allow the mountant to harden before being transferred to 4°C until imaging.

For NeuN staining, selected sections were rinsed in DPBS followed by washes in PBS with Triton. Sections were then incubated in a PBS with Triton solution containing donkey serum for 2 hours at room temperature. Sections were incubated overnight at 4°C in anti-NeuN antibody (Millipore Sigma, ABN78) at a 1:200 ratio. Following this incubation in primary antibody, sections were again washed in PBS with Triton and then incubated at room temperature in Donkey anti-Rabbit IgG (H+L) Highly Cross-Adsorbed Secondary Antibody, Alexa Fluor™ Plus 488 (Invitrogen, A32790) for 60 minutes at a 1:1000 ratio. Sections were then rinsed in PBS with Triton, then DPBS before mounting on slides. Once dry, sections were coverslipped using ProLong™ Gold Antifade Mountant with DNA Stain DAPI (ThermoFisher Scientific, P36935). Slides were stored at room temperature in dark conditions overnight to allow the mountant to harden before being transferred to 4°C until imaging.

### Imaging

Slides stored at 4°C were allowed to equilibrate to room temperature before imaging. Slides were imaged using an inverted microscope (Nikon ECLIPSE Ti2) with a Nikon Plan Fluor 10X Ph1 NA=0.30 Air objective using Nikon Elements software. Large area image acquisition was used to image the entirety of each brain section. Imaging was performed at Northwestern University’s Center for Advanced Microscopy.

### Imaging Analysis

Images were analyzed using FIJI software (version 2.16.0/1.54p). For analysis of images in the tyrosine hydroxylase (TH) staining cohort, red and blue channels were split, and red channel images were processed for analysis. For image processing, contrast was enhanced using 0.35% with normalization enabled. The image was then converted to 8-bit and Shanbhag Thresholding was applied. The resulting image was made binary, and watershed was applied to separate adjacent particles. FIJI’s Region of interest (ROI) manager was used to outline the left and right SNc and left and right VTA. FIJI’s “analyze particles” function was then used to count the number of particles in each region of interest with size set to 50-2000 square microns and circularity set at 0-1. Left and right SNc were combined for a single SNc value per section, as were the left and right VTA. ROI was defined by author D’Alessandro and sizing was kept consistent across sections. Two technical replicates were used for each biological replicate, and the technical replicates for each region (SNc and VTA) were averaged.

For analysis of images in the neuronal nuclei cohort, green and blue channels were split, and green channel images were processed for analysis. For image processing, the image was then converted to 8-bit and Moments Thresholding was applied. The resulting image was made binary, and watershed was applied to separate adjacent particles. ROI manager was used to outline the left and right SNc and left and right VTA. FIJI’s “analyze particles” function was then used to count the number of particles in each region of interest with size set to 50-30000 square microns and circularity set at 0.4-1. Left and right SNc were combined for a single SNc value per section, as were the left and right VTA. ROI was defined by author D’Alessandro and sizing was kept consistent across sections. Two technical replicates were used for each biological replicate, and the technical replicates for each region (SNc and VTA) were averaged.

For both tyrosine hydroxylase and neuronal nuclei groups, DAT-cre mouse cell counts for SNc and VTA were averaged across biological replicates. Using Excel, the average of the two technical replicates for each biological replicate in the MCI-Park and MCI-Park + MitoLbNOX LSL groups were calculated as a percentage of average DAT-Cre mouse expression. GraphPad Prism was then used to calculate p-values between groups.

For the representative images shown in Figure 4, contrast was enhanced using 0.35% with normalization enabled.

## Data Analysis

Data analysis was performed using GraphPad Prism (version 10.4.2). For metabolite ratios, peak intensities were compared within each sample. For the calculation of p-values, t-test with Welch’s correction was used. Multiple unpaired t-tests were used to calculate p-value for rotarod experiments. Grubbs’ test was applied to metabolomics and immunofluorescence data to determine any statistically significant outliers, which were removed. For survival analysis, log-rank (Mantel-Cox) test was used to determine p-values.

### Data Visualization

Figure 1a was created with BioRender.com. GraphPad Prism and Excel were used for data visualization.

## Acknowledgements

The authors would like to thank the Transgenic and Targeted Mutagenesis Laboratory, the Center for Advanced Microscopy, and the Behavioral Phenotyping Core at Northwestern University. We want to thank Dr. Craig Weiss and Jason Zysk from the Behavioral Phenotyping Core for their assistance. Imaging work was performed at the Northwestern University Center for Advanced Microscopy generously supported by NCI CCSG P30-CA060553 awarded to the Robert H. Lurie Comprehensive Cancer Center. We would also like to thank the staff at the Center for Comparative Medicine at Northwestern University. Additionally, we thank Dr. José López-Barneo and Dr. Lin Gao for providing our lab access to Ndufs2 floxed mice.

---

**Fig S1.**
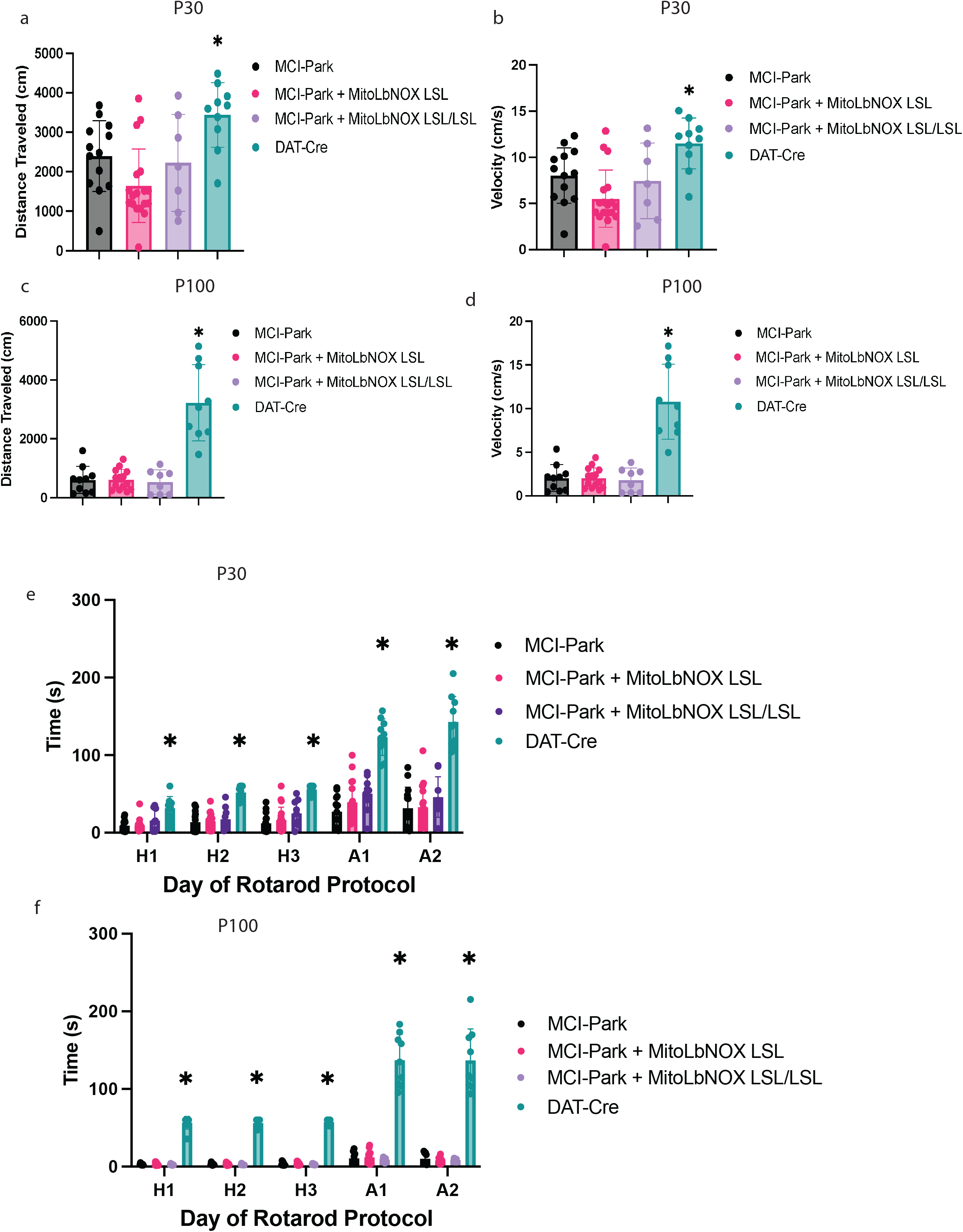
MitoLbNOX does not alter the behavioral phenotype of MCI-Park mice at P30 or P100. a) MitoLbNOX expression in MCI-Park mice does not restore open field distance traveled at P30 to the distance traveled in DAT-Cre mice (n=7-17 mice, Welch’s t-test p<0.05 for DAT-Cre mice compared to each other group, not significant for comparisons between MCI-Park, MCI-Park + MitoLbNOX, and MCI-Park + MitoLbNOX LSL/LSL). b) MitoLbNOX expression in MCI-Park mice does not restore open field velocity traveled at P30 to the velocity traveled in DAT-Cre mice (n=7-17 mice, Welch’s t-test p<0.05 for DAT-Cre mice compared to each other group, not significant for comparisons between MCI-Park, MCI-Park + MitoLbNOX, and MCI-Park + MitoLbNOX LSL/LSL). c) MitoLbNOX expression in MCI-Park mice does not restore open field distance traveled at P100 to the distance traveled in DAT-Cre mice (n=8-14 mice, Welch’s t-test p=0.0002 for DAT-Cre mice compared to each other group, not significant for comparisons between MCI-Park, MCI-Park + MitoLbNOX, and MCI-Park + MitoLbNOX LSL/LSL). d) MitoLbNOX expression in MCI-Park mice does not restore open field velocity traveled at P100 to the velocity traveled in DAT-Cre mice (n=8-14 mice, Welch’s t-test p=0.0002 for DAT-Cre mice compared to each other group, not significant for comparisons between MCI-Park, MCI-Park + MitoLbNOX, and MCI-Park + MitoLbNOX LSL/LSL). e) MitoLbNOX expression in MCI-Park mice does not restore rotarod performance at P30 to the performance levels of DAT-Cre mice (n=9-18 mice, multiple unpaired t-tests p<0.05 for DAT-Cre mice compared to each other group on all days, not significant for comparisons between MCI-Park, MCI-Park + MitoLbNOX, and MCI-Park + MitoLbNOX LSL/LSL on any day). f) MitoLbNOX expression in MCI-Park mice does not restore rotarod performance at P100 to the performance levels of DAT-Cre mice (n=7-13 mice, multiple unpaired t-tests p<0.000001 for DAT-Cre mice compared to each other group on all days, not significant for comparisons between MCI-Park, MCI-Park + MitoLbNOX, and MCI-Park + MitoLbNOX LSL/LSL on any day).

**Fig S2.**
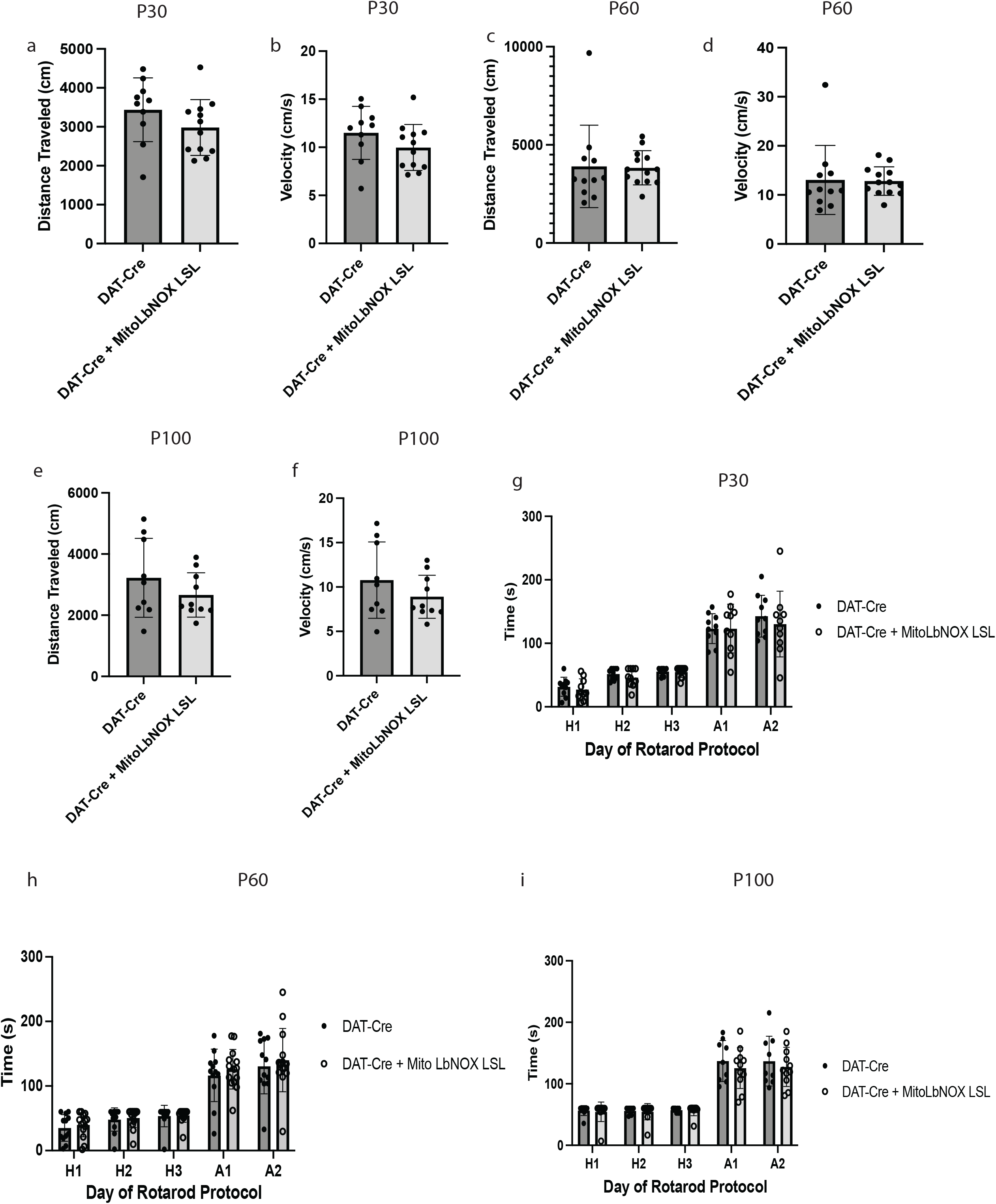
MitoLbNOX expression in DAT-Cre positive mice does not affect its motor phenotype. a) MitoLbNOX expression does not alter open field distance traveled at P30 compared to DAT-Cre control mice (n=10-12 mice, Welch’s t-test not significant). b) MitoLbNOX expression does not alter open field velocity traveled at P30 compared to DAT-Cre control mice (n=10-12 mice, Welch’s t-test not significant). c) MitoLbNOX expression does not alter open field distance traveled at P60 compared to DAT-Cre control mice (n=11-13 mice, Welch’s t-test not significant). d) MitoLbNOX expression does not alter open field velocity traveled at P60 compared to DAT-Cre control mice (n=11-13 mice, Welch’s t-test not significant). e) MitoLbNOX expression does not alter open field distance traveled at P100 compared to DAT-Cre control mice (n=9-10 mice, Welch’s t-test not significant). f) MitoLbNOX expression does not alter open field velocity traveled at P100 compared to DAT-Cre control mice (n=9-10 mice, Welch’s t-test not significant). g) MitoLbNOX expression does not alter rotarod performance at P30 compared to DAT-Cre control mice (n=10 mice, Welch’s t-test not significant). h) MitoLbNOX expression does not alter rotarod performance at P60 compared to DAT-Cre control mice (n=12-14 mice, Welch’s t-test not significant). i) MitoLbNOX expression does not alter rotarod performance at P100 compared to DAT-Cre control mice (n=9-11 mice, Welch’s t-test not significant).

